# Genomes of two indigenous clams *Anomalocardia flexuosa* (Linnaeus, 1767) and *Meretrix petechialis* (Lamarck, 1818)

**DOI:** 10.1101/2024.05.03.592324

**Authors:** Sean Tsz Sum Law, Wenyan Nong, Ming Fung Franco Au, Leni Hiu Tong Cheung, Cheryl Wood Yee Shum, Shing Yip Lee, Siu Gin Cheung, Jerome Ho Lam Hui

**Author notes:** contributed equally (Sean Tsz Sum Law, Wenyan Nong).

## Abstract

Clam digging has a long history in Hong Kong, but unregulated clam digging activities depletes clam populations and threatens the ecosystem. Population genomics is useful to unravel the connectivity of clams at different geographical locations and to provide necessary conservation measures; and yet, only limited number of clams in Hong Kong have genomic resources. Here, we present chromosomal-level genome assemblies for two clams commonly found in Hong Kong, *Anomalocardia flexuosa* and *Meretrix petechialis*, using a combination of PacBio HiFi and Omni-C reads. We assembled each genome (∼1.04-1.09 Gb) into 19 pseudochromosomes with high sequence continuity (scaffold N50 = 58.5 Mb and 53.5 Mb) and high completeness (BUSCO scores 94.4% and 95.7%). A total of 20,881 and 20,084 gene models were also predicted for *A. flexuosa* and *M. petechialis* respectively using transcriptomes generated in this study. The two new genomic resources established in this study will be useful for further study of the biology, ecology, and evolution of clams, as well as setting up a foundation for evidence-informed decision making in conservation measures and implementation.

## Background and Summary

Clams refer to the common name for several kinds of bivalve molluscs. The Veneridae family contains more than 700 described living species of bivalves or clams, and most of them are edible and exploited as food in different cultures around the world, including America, Asia and Europe (Huber, 2010)^1^. Clam digging activities, which refer to harvesting clams from below the surface of tidal sand or mud flats, also has long history in many places including Hong Kong. In the last century, clam digging in Hong Kong were mainly confined to villagers or recreational collection using hand tools on beaches during low tides for consumption or as a source of income. Nevertheless, clam digging activities have grown increasingly popular in recent years which threatens the clam populations and disturbs benthic biodiversity in some areas (Griffiths et al., 2006; So et al., 2021)^2,3^. Unlike many other places where sustainable clam digging practices, such as limiting the number of clams taken and/or temporary closure of clamming sites, Hong Kong does not have her own practices in the meantime due to the lack of information on the population structure of clams. Among the common clams that can be found in Hong Kong, such as that of *Anomalocardia* and *Meretrix* species which are the two frequently collected genera by local clam-diggers (So et al., 2021)^3^, genomic resources are currently lacking which hinders our understanding of their connectivity at different geographical locations.

Here, utilizing PacBio HiFi long reads and Omni-C sequencing data, we present two chromosomal-level genomes of common clams in Hong Kong, *Anomalocardia flexuosa* and *Meretrix petechialis*. Together with transcriptome data from various tissues, we produce high-quality predicted gene models for the two clam species. These genome assemblies and transcriptome data provide valuable genomic resources for the understanding of genetic diversity and connectivity for future population genomics research in view of conserving local clam species and assessing the sustainability of clam digging activities.

## Methods

### Sample collection and High molecular weight DNA extraction

*A. flexuosa* and *M. petechialis* samples were collected in Shui Hau, Lantau Island, Hong Kong (22°13’14.2”N 113°55’09.0”E) on 6^th^ July, 2023 and Yi O, Lantau Island, Hong Kong (22°13’58.4”N 113°51’02.0”E), on 28^th^ August, 2022, respectively. Approximately 300 mg adductor muscle was used for high molecular weight (HMW) DNA extraction for both *A. flexuosa* and *M. petechialis*. For *A. flexuosa*, the tissue was first ground into powder with liquid nitrogen, from which HMW DNA was isolated by NucleoBond HMW DNA kit (Macherey-Nagel), following the manufacturer’s protocol. For *M. petechialis*, HMW DNA was extracted using MagAttract HMW DNA Kit (Qiagen), following the manufacturer’s instructions. The DNA samples were eluted with 120 µL of elution buffer (PacBio Cat. No. 101-633-500) and were subjected to quality check by the Qubit® Fluorometer, NanoDrop One Spectrophotometer, and overnight pulse-field gel electrophoresis.

### PacBio library preparation and long-read sequencing

Prior to library preparation, approximately 5 µg of HMW DNA isolated from *A. flexuosa* and *M. petechialis* in 120µL of elution buffer were transferred to a g-tube (Covaris Cat. No. 520079) for DNA shearing with 6 passes of centrifugation at 1,990 x *g* for 2 min. The fragment size of sheared DNA samples was assessed with overnight pulse-field gel electrophoresis. A SMRTbell library was constructed for both samples using the SMRTbell® prep kit 3.0 (PacBio Cat. No. 102-141-700), following the manufacturer’s instructions. Qubit® Fluorometer and overnight pulse-field gel electrophoresis were used to examine the quantity and quality of the SMRTbell libraries. Subsequently, the Sequel®II binding kit 3.2 (PacBio Cat. No. 102-194-100) was used for the final library preparation with primer annealing, polymerase binding and the addition of internal DNA control. The two libraries were loaded at an on-plate concentration of 50-90 pM with diffusion loading mode. The sequencing was performed on the PacBio Sequel IIe system for a 30-hour movie to generate HiFi reads for each sample. One SMRT cell was used for sequencing for *A. flexuosa* and *M. petechialis*, respectively. Finally, 21.83 Gb and 30.58 Gb of Hifi reads were obtained for *A. flexuosa* and *M. petechialis* with average lengths of 8,017 bp and 10,729 bp and data coverages of 20X and 29X, respectively (Table 1).

### Omni-C library preparation and sequencing

An Omni-C library was prepared for *A. flexuosa* and *M. petechialis*, respectively, using the Dovetail® Omni-C® Library Preparation Kit (Dovetail Cat. No. 21005), following the manufacturer’s instructions. Approximately 50 mg of flash-freezing powered tissue was used for crosslinking with the addition of formaldehyde in 1 mL 1X PBS for each sample, followed by nuclease digestion. The lysate samples were assessed by Qubit® Fluorometer and TapeStation D5000 ScreenTape and were proceeded with the library preparation protocol. After the final quality check with Qubit® Fluorometer and TapeStation D5000 ScreenTape, the Omni-C libraries were sent to Novogene Co. Ltd for sequencing on an Illumina HiSeq-PE150 platform, from which 60.4 Gb and 56.6 Gb Omni-C data were generated for *A. flexuosa* and *M. petechialis*, respectively (Table 1).

### Transcriptome sequencing

Total RNA was isolated from various tissues including foot and adductor muscle, mantle, digestive gland, gill and gonad for *A. flexuosa* and foot, digestive gland, gill and gonad for *M. petechialis*, using the mirVana™ miRNA Isolation Kit (Ambion), following the manufacturer’s protocol respectively. The RNA samples were subjected to quality control using NanoDrop One Spectrophotometer, and gel electrophoresis. The qualified samples were sent to Novogene Co. Ltd for polyA selected RNA sequencing library construction and 150 bp paired-end sequencing. A total of 31.7 Gb and 23.1 Gb transcriptome data were obtained from different tissue types of *A. flexuosa* and *M. petechialis*, respectively (Table 1).

### Genome assembly and Gene model prediction

*De novo* genome assemblies of *A. flexuosa* and *M. petechialis* were first proceeded with Hifiasm (Cheng et al., 2021)^4^ and then were processed with searching against the NT database with BLAST to remove possible contaminations using BlobTools (v1.1.1) (Laetsch & Blaxter, 2017)^5^. Subsequently, haplotypic duplications were removed according to the depth of HiFi reads using “purge_dups” (Guan et al., 2020)^6^. Proximity ligation data from Omni-C were used to scaffold the assembly using YaHS (Zhou et al., 2022)^7^ and manual checking using Juicebox (v1.1)^8^. The genomes were soft-masked by redmask (v0.0.2) (https://github.com/nextgenusfs/redmask) (Girgis et al., 2015)^9^. The final genome assemblies of *A. flexuosa* and *M. petechialis* were 1.09 Gb and 1.04 Gb in size with 95.43% and 99.27% of the sequenced anchored into 19 chromosomes, respectively, which correspond to the kartotype (2n = 38) of *Anomalocardia* and *Meretrix* species (Figure 1B-C; Table 2-3) (Lavander et al., 2017; Park et al., 2011)^10,11^. Both *A. flexuosa* and *M. petechialis* genomes were not only of high continuity, with scaffold N50 of 58.5 Mb and 53.5 Mb in 9 scaffolds, but also of high completeness after being assessed with Benchmarking Universal Single-Copy Orthologs (BUSCO, v5.5.0) using the “metazo_odb10” dataset (Manni et al., 2021)^12^, which resulted in BUSCO scores of 94.4% and 95.7%, respectively (Figure 1B; Table 2).

**Figure 1.**
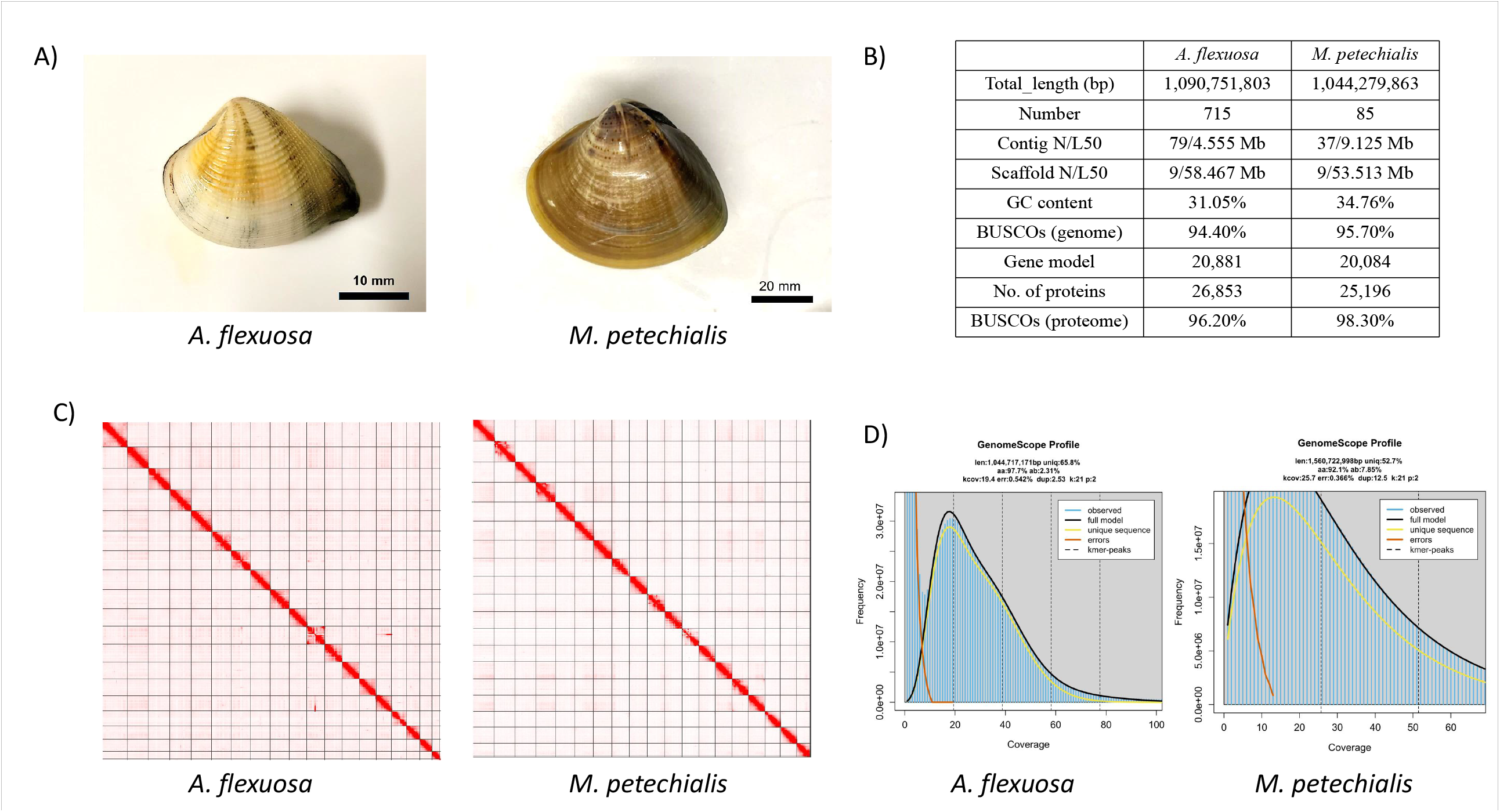
A) Pictures of *A. flexuosa* (left) and *M. petechialis* (right); B) Statistics of the genome assembly generated in this study; C) Hi-C contact map of the assembly *A. flexuosa* (left) and *M. petechialis* (right); D) Estimated genome size (*K*-mer = 21) of *A. flexuosa* (left) and *M. petechialis* (right).

For gene model prediction, RNA sequencing data were first processed using Trimmomatic (v0.39) (Bolger, Lohse & Usadel 2014)^13^ with parameters “TruSeq3-PE.fa:2:30:10 SLIDINGWINDOW:4:5 LEADING:5 TRAILING:5 MINLEN:25” and kraken2 (v2. 0.8 with kraken2 database k2_standard_20210517)^14^ to remove the low quality and contaminated reads, and then aligned to the repeat soft-masked genome using Hisat2^15^ to generate the bam file. A total of 389,399 Mollusca reference protein sequences were downloaded from NCBI on 25 Mar 2024 as protein hits, along with the RNA bam file, to perform genome annotation using Braker (v3.0.8)^16^ with default parameters. These data collectively generated 20,881 and 20,084 gene models for *A. flexuosa* and *M. petechialis*, comprising 26,853 and 25,196 predicted protein-coding genes with average lengths of 580 and 607 amino acids, respectively (Figure 1B; Table 2). The completeness of proteomes were also evaluated with BUSCO “metazo_odb10” dataset (Manni et al., 2021)^12^, reporting BUSCO scores of 96.2% and 98.3%, respectively (Figure 1B; Table2).

### Repetitive elements annotation

Transposable elements (TEs) of the two genome assemblies were annotated as previously described (Baril et al, 2024)^17^ using the automated Earl Grey TE annotation pipeline (version 1.2, https://github.com/TobyBaril/EarlGrey) with “-r eukarya” to search the initial mask of known elements and other default parameters. Briefly, this pipeline first identified known TEs from Dfam with RBRM (release 3.2) and RepBase (v20181026). *De novo* TEs were then identified, and consensus boundaries were extended using an automated “BLAST, Extract, Extend’ process with 5 iterations and 1000 flanking bases added in each round. Redundant sequences were removed from the consensus library before the genome assembly was annotated with the combined known and *de novo* TE libraries. Overlap and defragment annotations were removed prior to final TE quantification. A total of 338.3 Mb and 427.2 Mb of repeat contents were annotated from the genomes of *A. flexuosa* and *M. petechialis*, which account for 31.05% and 40.85% of the assembly, respectively (Figure 2; Table 4). Of the classified TEs, LINE, DNA, and Rolling Circle contribute to the major proportions (Figure 2), which are listed in Table 4.

**Figure 2.**
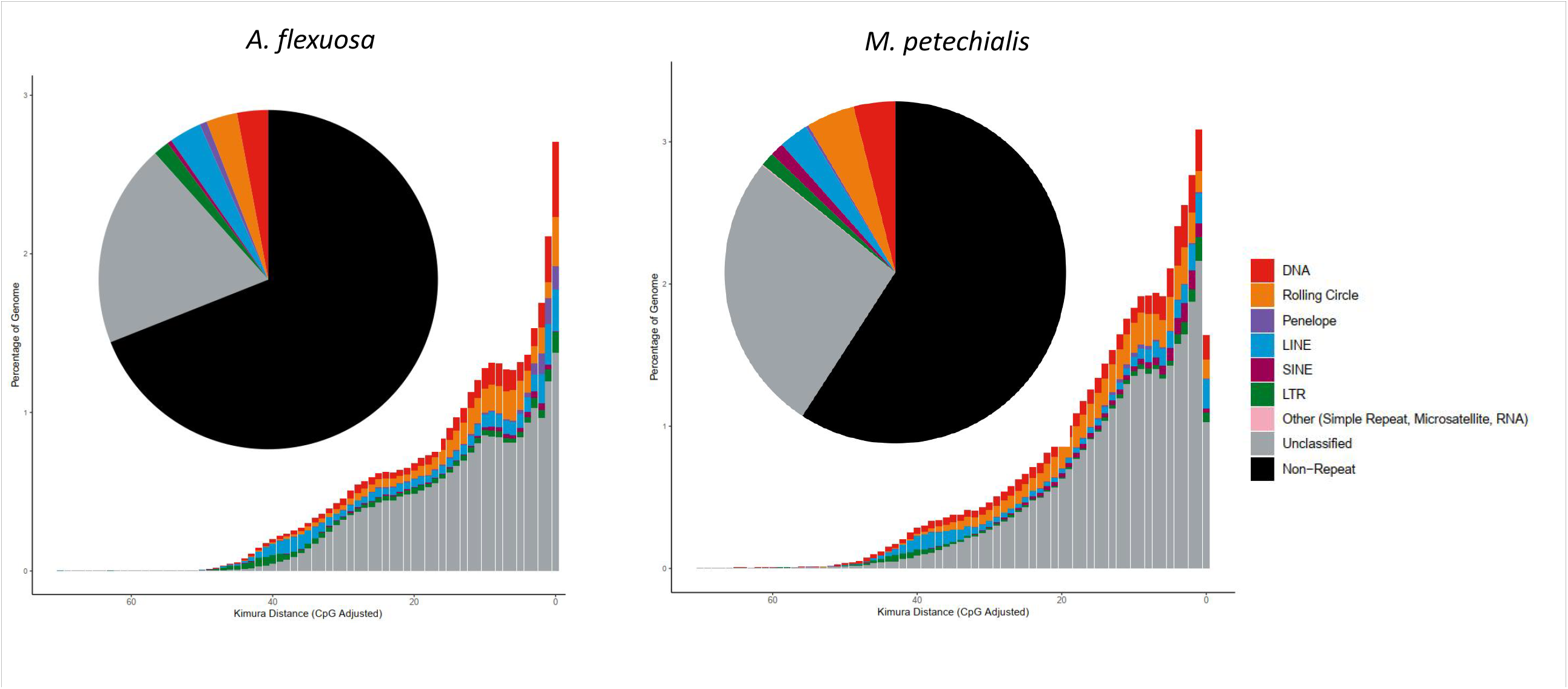
Repeat content of *A. flexuosa* (left) and *M. petechialis* (right).

### Syntenic analyses

Macrosynteny analysis revealed a 1-to-1 pair relationship between the 19 pseudochromsomes of *A. flexuosa* and *M. petechialis* using JCVI utility libraries ^18^(Figure 3), showing a conserved chromosome architecture among the two species.

**Figure 3.**
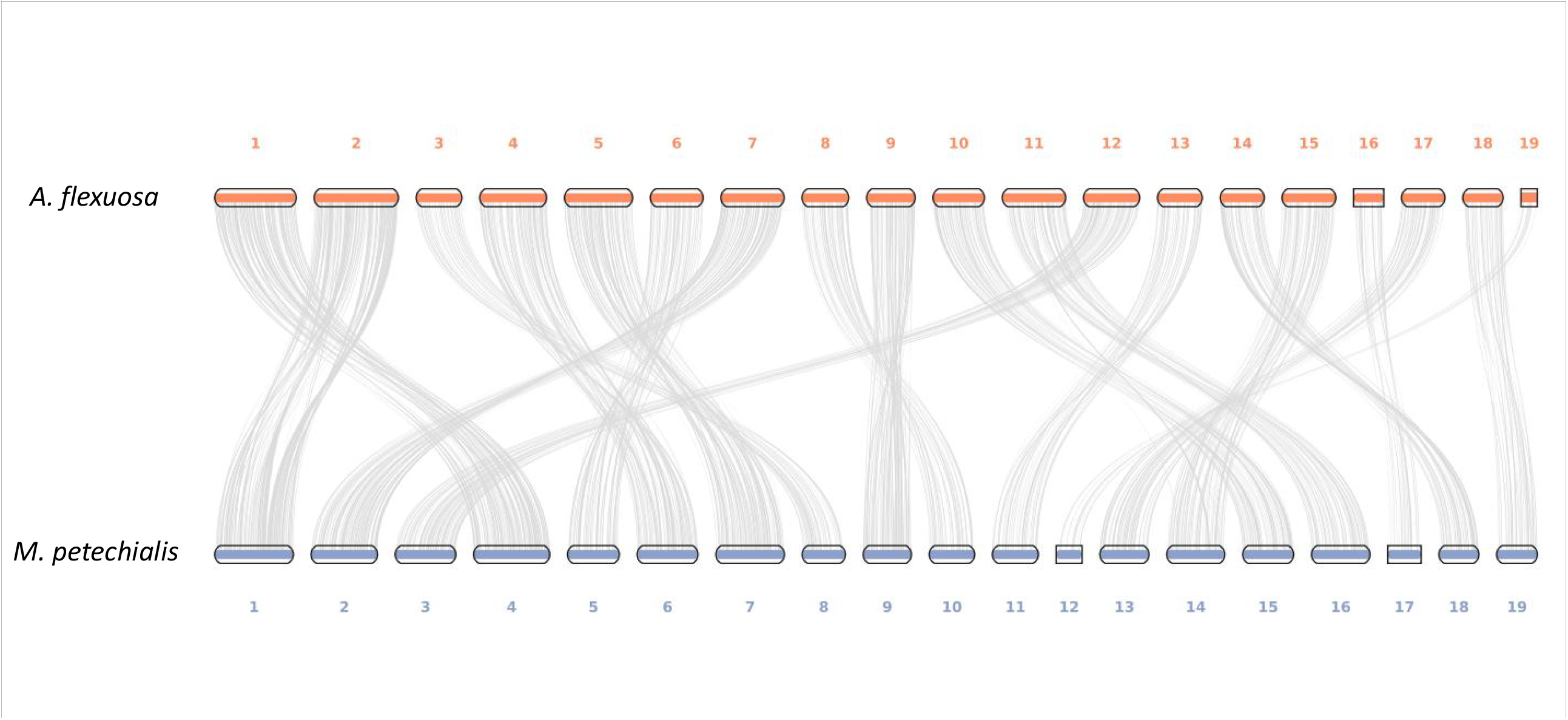
Macrosynteny plot of the 19 pseudochromosomes between *A. flexuosa* and *M. petechialis*.

**Figure 4.**
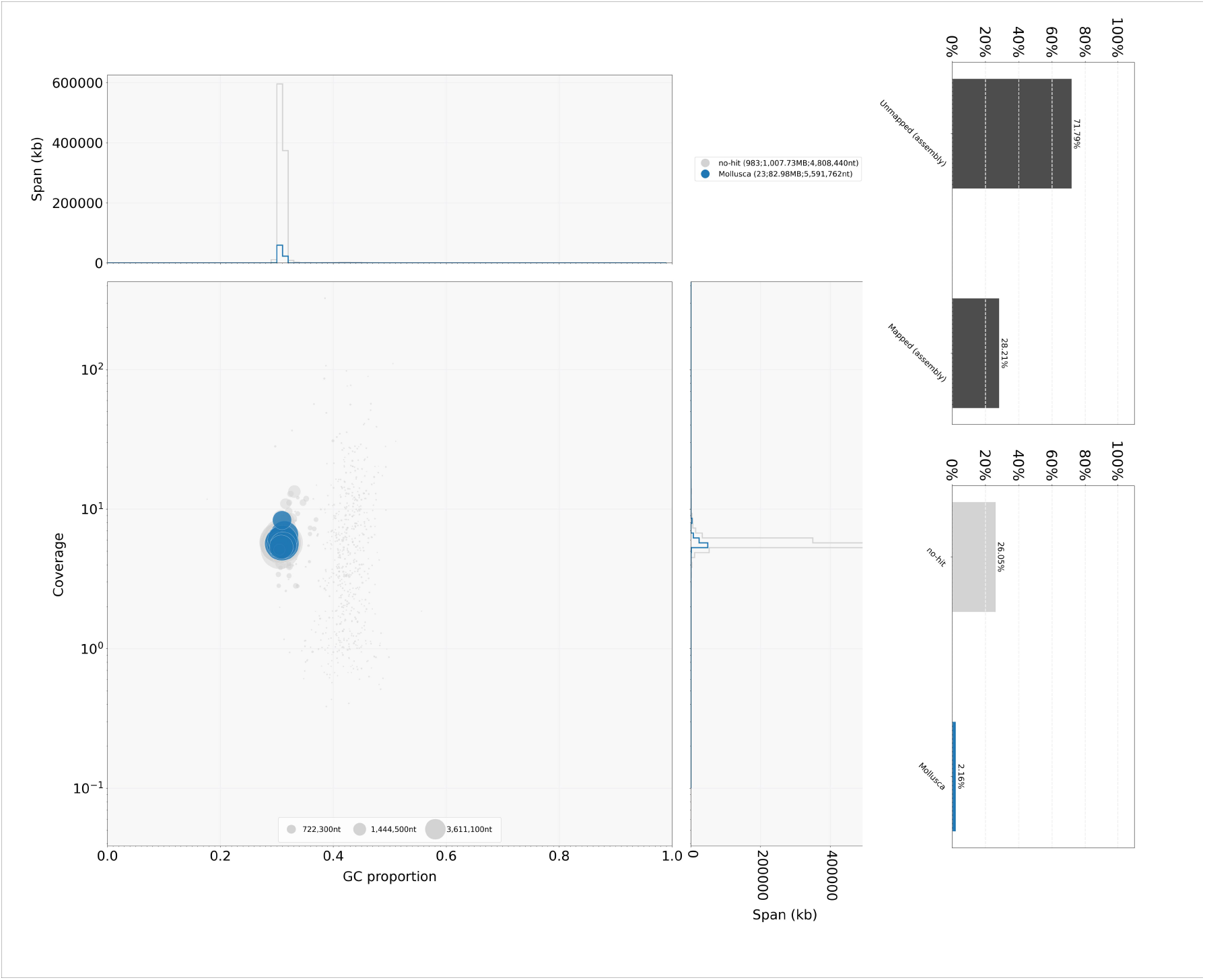
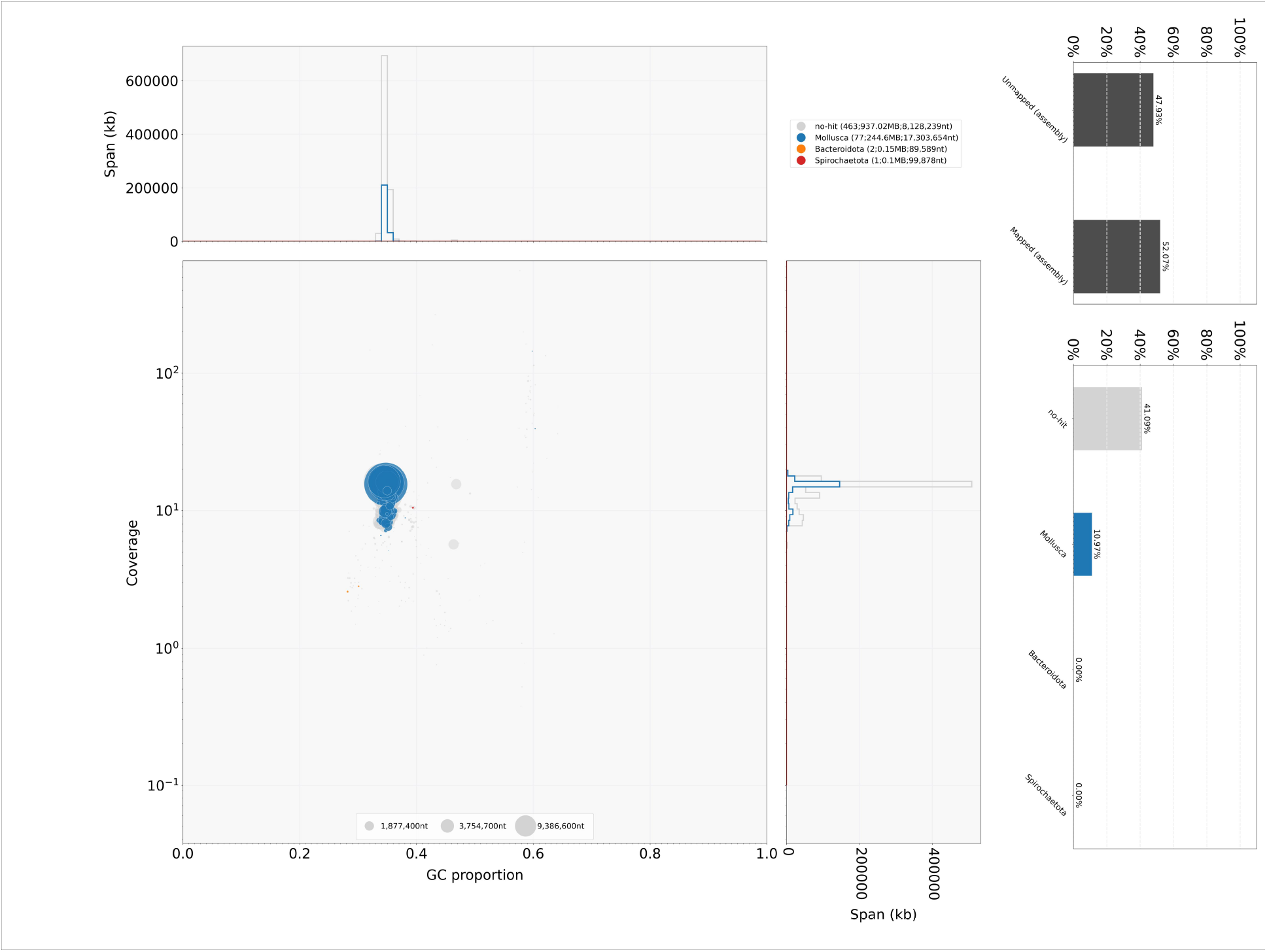
Genome assembly quality control (QC) and contaminants detection for *A. flexuosa* **(a)** and *M. petechialis* **(b)**.

## Data Records

The raw reads generated in this study, including Transcriptome, Omni-C and PacBio HiFi data, have been deposited in the NCBI database under the BioProject accession number PRJNA1100776 (https://www.ncbi.nlm.nih.gov/bioproject/PRJNA1100776) and PRJNA1100773 (https://www.ncbi.nlm.nih.gov/bioproject/PRJNA1100773) for *A. flexuosa* and *M. petechialis*, respectively. The genome, genome and repeat annotation files have been deposited and are publicly available in Figshare (https://figshare.com/s/88cafb78affd49b5b3ee).

## Technical Validation

The pseudochromosomes of the final assemblies were validated by inspecting the Omni-C contact maps using Juicer tools (version 1.22.01) (Durand et al., 2016)^8^. Briefly, Omni-C reads were mapped and aligned by BWA with parameters “mem -5SP -T0”, the parsing module of the pairtools pipeline was used to find ligation junctions with parameters “--min-mapq 40 -- walks-policy 5unique --max-inter-align-gap 30 --nproc-in 8 --nproc-out 8”. The parsed pairs were then sorted using pairtools sort with default parameters, PCR duplicate pairs were removed using pairtools dedup with parameters “--nproc-in 8 --nproc-out 8 --mark-dups”, the pairs file was split using pairtools split with default parameters and used to generate the contact matrix using juicertools and Juicebox (v1.11.08)^8^. Regarding the genome characteristics of the assembly, the k-mer count and histogram were generated at k = 21 from Omni-C reads using Jellyfish (v2.3.0) (Marçais & Kingsford, 2011)^19^ with the parameters “count -C -m 21 -s 1000000000 -t 10”, and the reads.histo was uploaded to GenomeScope to estimate genome heterozygosity, repeat content and size using default parameters (v2.0) (http://qb.cshl.edu/genomescope/genomescope2.0/) (Ranallo-Benavidez et al., 2020)^20^. The resulting GenomeScope plots can be found in Figure 1D.

Code availability N/A.

## Acknowledgements

This work was supported by Lantau Conservation Fund (RE-2020-39), Hong Kong Research Grant Council Collaborative Research Fund (C4015-20EF), Innovation Technology Fund of Innovation Technology Commission: Funding Support to State Key Laboratory of Agrobiotechnology, and Direct Grant of The Chinese University of Hong Kong (4053618).

## Contributions

S.Y.L, S.G.C and J.H.L.H conceived and supervised the study; S.T.S.L, M.F.F.A, L.H.T.C and C.W.Y.S carried out sample collection; S.T.S.L and W.N. performed data curation on the analysis; J.H.L.H., S.T.S.L. and W.N. wrote the initial manuscript; all authors revised and contributed to the final version of the manuscript.

## Corresponding author

Correspondence to Shing Yip Lee, Siu Gin Cheung, Jerome Ho Lam Hui.

## Tables and Figures

**Table 1**. Genome and transcriptome sequencing data

**Table 2**. Genome statistics.

**Table 3**. Psuedochromosome information.

**Table 4**. Repeat content summary.

